# Mucosal and systemic neutralizing antibodies to norovirus and rotavirus by oral immunization with recombinant rotavirus in infant mice

**DOI:** 10.1101/2022.09.01.505917

**Authors:** Takahiro Kawagishi, Liliana Sánchez-Tacuba, Ningguo Feng, Veronica P. Costantini, Ming Tan, Xi Jiang, Kim Y. Green, Jan Vinjé, Siyuan Ding, Harry B. Greenberg

**Affiliations:** Department of Medicine, Division of Gastroenterology and Hepatology, Stanford University School of Medicine, Stanford, CA; Department of Microbiology and Immunology, Stanford University School of Medicine, Stanford, CA; VA Palo Alto Health Care System, Department of Veterans Affairs, Palo Alto, CA; National Calicivirus Laboratory, Centers for Disease Control and Prevention, Atlanta, GA; Divison of Infectious Diseases, Cincinnati Children’s Hospital Medical Center, Cincinnati, OH; Department of Pediatrics, University of Cincinnati College of Medicine, Cincinnati, OH; Laboratory of Infectious Disease, National Institute of Allergy and Infectious Diseases, National Institutes of Health, Bethesda, MD; Department of Molecular Microbiology, Washington University School of Medicine, St. Louis, MO

**Keywords:** Mucosal vaccination, enteric viral vector, rotavirus, norovirus, enteric pathogens

## Abstract

Rotaviruses (RVs) preferentially replicate in the small intestine, frequently cause severe diarrheal disease, and following enteric infection generally induce variable levels of protective systemic and mucosal immune responses in humans and other animals. Rhesus rotavirus (RRV) is a simian RV that was previously used as a human RV vaccine and has been extensively studied in mice. Although RRV replicates poorly in the suckling mouse intestine, infection induces a robust and protective antibody response. The recent availability of plasmid-based RV reverse genetics systems has enabled the generation of recombinant RVs expressing foreign proteins. However, recombinant RVs have not yet been experimentally tested as potential vaccine vectors to immunize against other gastrointestinal pathogens *in vivo*. This is a missed opportunity because several live-attenuated RV vaccines are already widely administered to infants and young children worldwide. To explore the feasibility of using RV as a dual vaccine vector, we rescued a replication-competent recombinant RRV harboring bicistronic gene segment 7 that encodes both the native RV NSP3 protein and a human norovirus (HuNoV) VP1 protein from the predominant genotype GII.4 (rRRV-HuNoV-VP1). The rRRV-HuNoV-VP1 expressed HuNoV VP1 in infected cells *in vitro* and importantly, elicited both systemic and local antibody responses to HuNoV following oral infection of suckling mice. Serum IgG and fecal IgA from infected suckling mice bound to and neutralized both RV and HuNoV. These findings have encouraging practical implications for the design of RV-based next-generation multivalent enteric vaccines to target HuNoV and other human enteric pathogens while providing immunity to RV.

**Significance statement:** Mucosal immunity is a key component of protection against many pathogens. Robust and effective mucosal immune responses are generally induced following infection with a replication-competent pathogen at a mucosal surface. Several studies have attempted to develop viral vector-based enteric mucosal vaccines; however, the most advanced of these are still in clinical development. Here, we successfully induced systemic and mucosal antibody responses against both rotavirus and norovirus following inoculation of a recombinant rotavirus expressing the human norovirus major capsid protein. These responses are likely to correlate with protective immunity. Live-attenuated rotavirus vaccines have already proven safe and effective worldwide. These findings confirm the potential utility of using rotaviruses as a dual enteric vaccine platform for other important human enteric pathogens.

## Introduction

Mucosal immunity plays a critical role in protecting against many pathogens in the respiratory and intestinal tracts. Live virus infections have generally triggered more robust and effective mucosal immune response than oral administration of inactivated viruses or target protein antigens because they are self-amplifying and can elicit cellular as well as humoral immunity (1-4). Several studies have attempted to utilize recombinant viruses as vaccine vectors to induce an immune response against enteric pathogens (5-8); however, the most advanced of such enteric vaccine vectors are still in early stages of clinical development.

Rotaviruses (RVs), the leading cause of acute gastroenteritis in infants, are a promising candidate for enteric vaccine vectors for several reasons. (A) RV preferentially replicates in the small intestine, distinguishing it from several other enteric viruses that can also infect systemically or the colon. (B) RV infection is acute, and the virus does not integrate into the host genome. (C) RV is highly immunogenic and induces both systemic and mucosal immune responses in infected animals and humans (9, 10). (D) Several live-attenuated human RV vaccines have been shown to be both safe and effective to use in very young children (e.g., RotaTeq [Merck] and Rotarix [GlaxoSmithKline]). Other effective live-attenuated RV vaccines (Rotasiil, Rotavac, LLR, and Rotavin-M1) are also licensed for use globally or primarily in their country of origin (11). (E) Following substantial public health efforts, RV vaccines are now widely available in many low- and middle-income countries, as well as the more developed countries, hence the administration of RV-based vaccines that included other heterologous antigens could potentially be piggybacked onto current RV immunization programs used globally. (F) The RV double-stranded RNA (dsRNA) genome is segmented in nature, permitting easy genetic manipulation. (G) With the insertion of heterologous antigens, RV replication can become attenuated and has been shown to have reduced pathogenicity (12, 13).

Since a plasmid-based reverse genetics system was established in 2017, several studies have reported the generation of recombinant RVs that express fluorescent and bioluminescent reporter proteins (GFP, RFP, luciferase, etc.) and exogenous nucleotide sequences (e.g., endoribonuclease Csy4 target sequence and sequences encoding the receptor binding domain of the SARS-CoV-2 spike protein) *in vitro* (12-22). To facilitate the assessment and development of RVs as potential enteric vaccine vectors, the capacity of recombinant RVs to induce an enteric immune response against other gastrointestinal (GI) pathogens needs to be evaluated in well characterized pre-clinical small animal models. Rhesus rotavirus (RRV) is a prototype laboratory strain of simian RV that efficiently replicates *in vitro* (23, 24). Although RRV does not replicate well in a murine model (25-27), it does induce both systemic and mucosal immune responses in infected mice (28). In addition, RRV itself and RRV-based RV vaccine candidates have previously been shown to be a highly immunogenic and protective in several previous human vaccine trials and were, for a time, licensed for use in children in the United States (29, 30).

Human norovirus (HuNoV) is a major cause of acute gastroenteritis in both young children and adults. Although B cells and human intestinal organoids support HuNoV replication (31, 32), there is not yet a widely available robust cell culture system for efficient HuNoV cultivation, which has impeded both the assessment of HuNoV immunity and vaccine development. The HuNoV virion consists of major capsid protein VP1 and minor capsid protein VP2 surrounding a positive-sense RNA genome (33-35). Exogenously expressed VP1 can form virus-like particles (VLPs) that are structurally and antigenically similar to HuNoV virions (36-38) and the parenteral administration of such VLPs provides some level of protective immunity to HuNoV in adults (39-41). Moreover, expression of the protruding or P domain of VP1 that bears the major antigenic sites of HuNoV can yield subunit “P particles” that can also induce immune responses (42, 43). Here, we demonstrate the induction of both systemic and mucosal antibody responses against HuNoV in suckling mice using a recombinant RRV expressing HuNoV VP1. Our data suggest that recombinant RVs represent a potentially effective small-intestine-targeted vaccination platform to express exogenous genes in the human intestine to protect people from other enteric pathogens such as HuNoV as well as RV.

## Results

### Generation of a recombinant RV expressing HuNoV VP1

To express HuNoV VP1 from RV, we first constructed a plasmid with a T7 promoter driving the expression of RV nonstructural protein 3 (NSP3), a “self-cleavage” peptide derived from Thosea asigna virus 2A (T2A), and the coding sequence of major capsid protein VP1 of GII.4 HuNoV Sydney strain into the RV gene segment 7 (**Figure S1A**). We confirmed the expression of the expected proteins from this construct design by immunostaining (**Figure S1B**) and western blot (**Figure S1C**) following transfection of this plasmid into BHK-T7 cells. A faint signal for the fusion protein (RV-NSP3-T2A-HuNoV-VP1) was also detected by both the HuNoV VP1- and RV NSP3-specific sera in the western blot, but the cleavage mediated by T2A to release NSP3 (36 kDa) and VP1 (∼58 kDa) appeared quite efficient (**Figure S1C**). Using an improved RV reverse genetics system previously described (12), we rescued a recombinant RRV harboring the HuNoV VP1 gene (rRRV-HuNoV-VP1). To confirm that the rRRV-HuNoV-VP1 actually harbored the modified gene segment 7, we extracted viral dsRNA from virions of rRRV-HuNoV-VP1 and compared the electrophoretic pattern of the dsRNA with that from wild-type rRRV. RNA-PAGE analysis showed that rRRV-HuNoV-VP1 passage 3 stock has an extra band between gene segment 1 (3,302 bp) and gene segment 2 (2,708 bp), which corresponds in size to the modified gene segment 7 size (2,752 bp). Of note, the rRRV-HuNoV-VP1 lost this extra band between gene segments 1 and 2 during serial passage (**Figure 1A**). This finding suggests that the modified gene segment 7 was incorporated into progeny virion, but that the resulting recombinant RV was not genetically stable during multiple passages in cell culture. Growth kinetics of rRRV-HuNoV-VP1 in MA104 cells showed that the virus replicated efficiently at multiplicity of infection (MOI) of 1 and 0.01 focus forming unit (FFU)/cell although the recombinant HuNoV-VP1 expressing virus titer was slightly lower than that of wild-type rRRV (**Figures 1B and 1C**). We next demonstrated that rRRV-HuNoV-VP1 expressed HuNoV VP1 in RV-infected cells by immunostaining (**Figure 1D**) and determined that the molecular size of the expressed HuNoV VP1 corresponded to HuNoV VLP by western blot assay (**Figure 1E**). Taken together, these data demonstrate that rRRV-HuNoV-VP1 harbors the modified RV gene segment 7 during viral replication and produces HuNoV VP1 in infected cells as designed. Because of the limited genetic stability of rRRV-HuNoV-VP1, we used passage 3 stocks in all the subsequent animal experiments described below.

**Figure 1.**
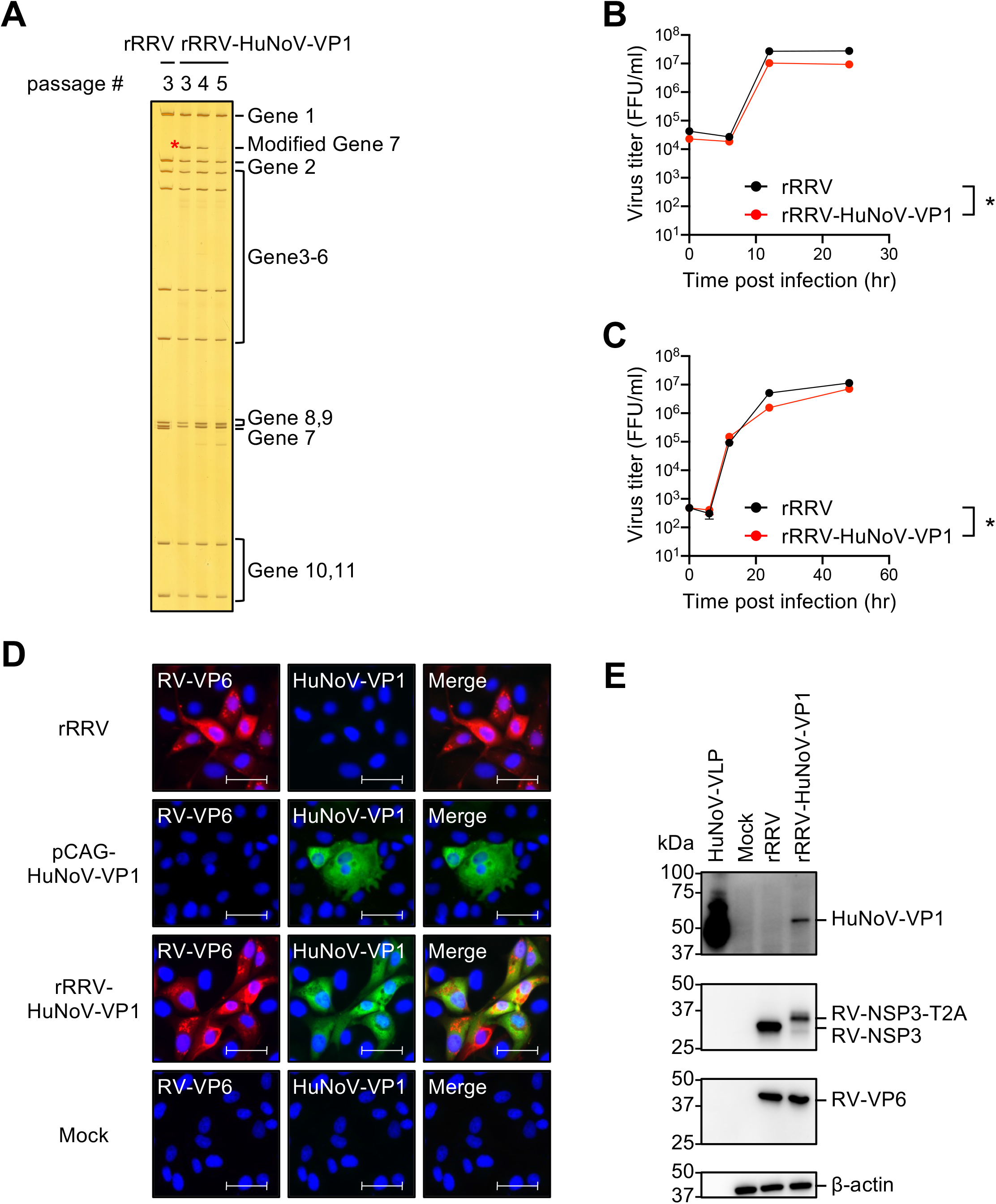
*In vitro* characterization of rRRV-HuNoV-VP1. (**A**) Electrophoretic analysis of viral dsRNA. Viral dsRNAs were purified from rRRV and different passages of rRRV-HuNoV-VP1, separated by RNA-PAGE, and visualized by silver staining. The modified gene segment 7 is indicated by a red asterisk. (**B and C**) Growth kinetics of rRRV and rRRV-HuNoV-VP1 in MA104 cells. MA104 cells were infected with either rRRV or rRRV-HuNoV-VP1 at an MOI of (**B**) 1 FFU/cell or (**C**) 0.01 FFU/cell. The samples were frozen at each indicated time point and the virus titers were determined by a focus forming unit (FFU) assay. Data are shown as mean with standard deviation. Statistical analysis was performed by two-way ANOVA. Statistical significance is indicated as *: P < 0.05. (**D**) Immunostaining analysis of protein expression by rRRV-HuNoV-VP1 in MA104 cells. MA104 cells were infected with the rRRV or rRRV-HuNoV-VP1 at MOI of 1 FFU/cell and fixed at 24 hours post infection, or the cells were transfected with pCAG-HuNoV-VP1 plasmid and fixed at 3 days post transfection. The cells were stained with antibodies specific to RV-VP6 (red), HuNoV-VP1 (green), and DAPI (blue). Scale bar: 50 μm. (**E**) Western blotting analysis of protein expression by rRRV-HuNoV-VP1 in MA104 cells. MA104 cells were infected with rRRV or rRRV-HuNoV-VP1 at an MOI of 4 FFU/cell and harvested at 12 hours post infection. The cells were lysed with Laemmli buffer and resolved by SDS-PAGE. Protein expression of HuNoV-VP1, RV-NSP3, RV-VP6, and β-actin was detected by the specific antibodies. The numbers show the molecular weights determined by the protein ladder.

### Effect of HuNoV VP1 gene insertion on RV replication and pathogenesis *in vivo*

Although RRV does not replicate robustly in the small intestine of neonatal mice, it causes diarrhea in pups at early time points following inoculation (26, 44). To assess the effect of HuNoV VP1 insertion on RRV replication *in vivo*, we inoculated immunocompetent 5-day-old 129sv pups with 3.9×10^5^ FFUs of rRRV-HuNoV-VP1 or the parental rRRV as a control and compared the diarrhea rate and fecal RV shedding over ten days. Like rRRV, rRRV-HuNoV-VP1 caused diarrhea at early times post-inoculation in 129sv pups, but neither virus was detectable by ELISA in the stools of the 129sv pups (**Figures 2A and 2B**). Previous studies demonstrated that RRV replicates better in immunodeficient *Stat1*^*-/-*^ pups than in immunocompetent pups, presumably because STAT1 serves as a vital mediator in the interferon-induced anti-RV signaling cascade (26, 27, 45). To test replication of rRRV-HuNoV-VP1 in suckling mice under less restrictive growth conditions, we examined diarrheal rates and fecal RV shedding in 5-day-old *Stat1*^*-/-*^ pups. Both rRRV and rRRV-HuNoV-VP1 caused diarrhea from 1 to 4 days post-infection (**Figure 2C**). We detected modest fecal RV shedding by rRRV and rRRV-HuNoV-VP1 from 1 to 7 days post-inoculation in the *Stat1*^*-/-*^ pups (**Figure 2D**). The amount of fecal RV shedding by rRRV-HuNoV-VP1 was not significantly different from that by rRRV (**Figure 2D**). These data suggest that rRRV-HuNoV-VP1 replicates to a similar extent as rRRV in both 129sv and *Stat1*^*-/-*^ mice and that both viruses replicated significantly more in the *Stat1*^*-/-*^ pups.

**Figure 2.**
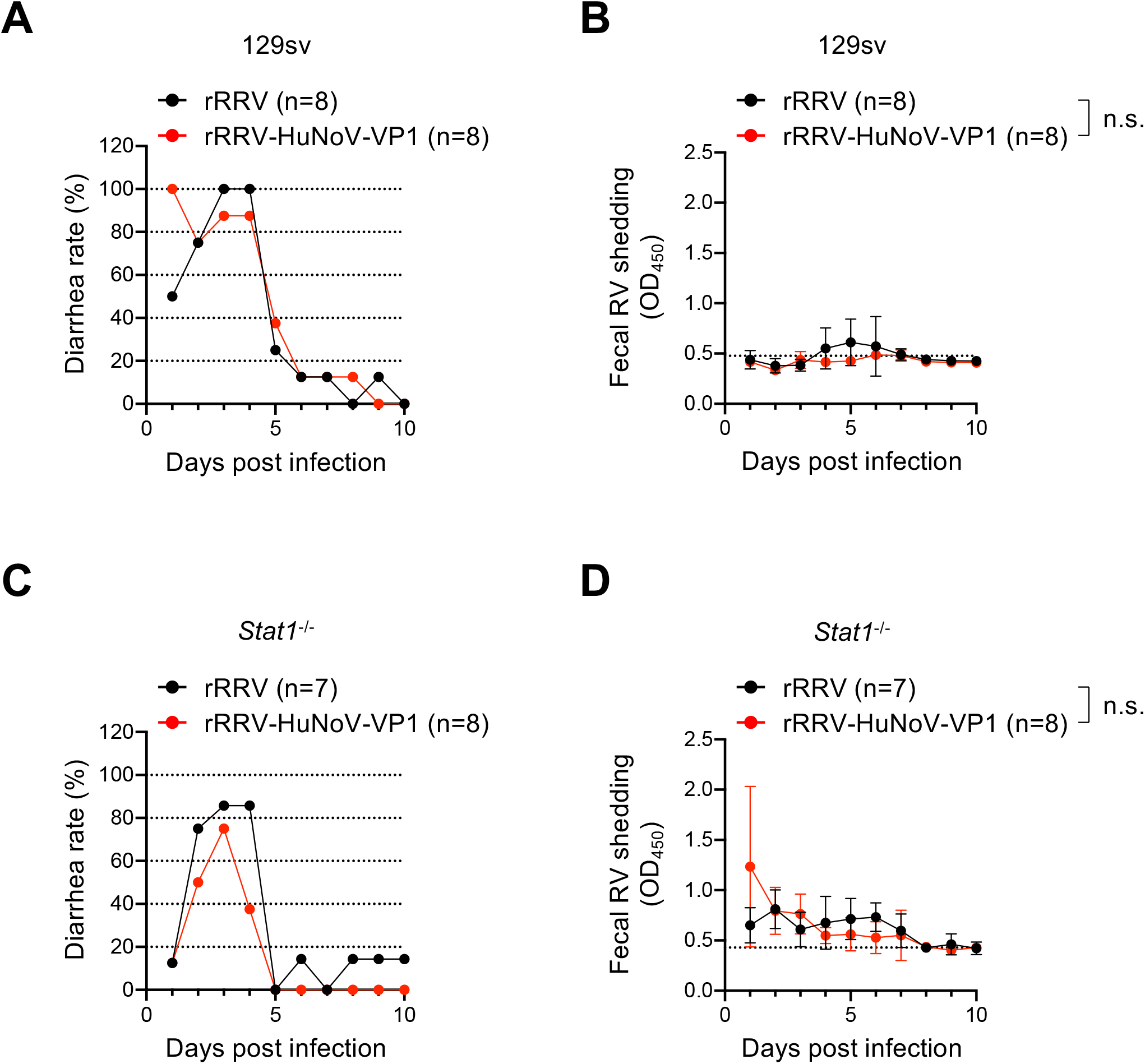
*In vivo* characterization of rRRV-HuNoV-VP1. (**A and C**) Diarrhea rate by rRRV and rRRV-HuNoV VP1 in (**A**) 129sv or (**C**) *Stat1*^*-/-*^ mice. Five-day-old 129sv pups were orally inoculated with rRRV (n=8) or rRRV-HuNoV-VP1 (n=8) (3.9×10^5^ FFU/pup), and five-day-old *Stat1*^*-/-*^ pups were orally inoculated with rRRV (n=7) or rRRV-HuNoV-VP1 (n=8) (3.9×10^5^ FFU/pup). Diarrhea was monitored daily following gentle abdominal palpation until 10 days post-infection. The daily percentage of diarrhea is shown. (**B and D**) Fecal shedding curve in (**B**) 129sv or (**D**) *Stat1*^*-/-*^ mice after rRRV or rRRV-HuNoV-VP1 inoculation. Fecal RV antigens in stools collected from 1 to 10 days post infection were measured by ELISA. Data are shown as mean scores of net OD_450_ with standard deviation. The dotted lines show background signals measured by stools from uninfected pups. Statistical analysis was performed with two-way ANOVA. Statistical significance is indicated as n.s.: not significant.

### Induction of serum IgG responses in 129sv and *Stat1*^*-/-*^ suckling mice following rRRV-HuNoV-VP1 infection

Next, we examined the ability of rRRV-HuNoV-VP1 to induce a systemic antibody response against HuNoV VP1 in the suckling mice. We orally inoculated 5-day-old 129sv and *Stat1*^*-/-*^ pups with 3.9×10^5^ FFUs of rRRV-HuNoV-VP1 or the wild-type rRRV and collected blood and stool specimens at 4-, 6-, and 8-weeks post inoculation (WPI). In the case the primary infection with rRRV-HuNoV-VP1 induced no or a weak immune response, we also provided a parenteral intraperitoneal (IP) booster immunization with the same RV dose (3.9×10^5^ FFU/pup) at 9 WPI and collected blood and stool specimens one week later (10 WPI) (**Figure 3A**). To examine if the mouse sera contained antibodies against HuNoV VP1, we first established a new immunostaining assay. We expressed either HuNoV VP1 or RV VP6, a broadly conserved RV antigen, in BHK-T7 cells, and stained for these proteins with the mouse sera collected at 4 WPI. Consistent with previous studies (28), sera from 129sv pups infected with either rRRV-HuNoV VP1 or rRRV group reacted robustly to the BHK-T7 cell expressed RV VP6. However, we did not observe a clear HuNoV VP1 staining signal with these sera (**Figures S2A and S2B**). The sera from an uninfected 129sv mouse served as a negative control and, as expected, failed to react with either RV VP6 or HuNoV VP1 (**Figure S2C**). Interestingly, we found that sera from nine of the eleven *Stat1*^*-/-*^ mice inoculated with rRRV-HuNoV-VP1 recognized both RV VP6 and HuNoV VP1 (**Figure S3A**). As expected, the sera from *Stat1*^*-/-*^ mice infected with rRRV only reacted with RV VP6, while the sera from an uninfected *Stat1*^*-/-*^ mouse did not react to either RV VP6 or HuNoV VP1 (**Figures S3B and S3C**). These data suggest that the sera from rRRV-HuNoV-VP1-infected *Stat1*^*-/-*^ mice reacted specifically to HuNoV VP1.

**Figure 3.**
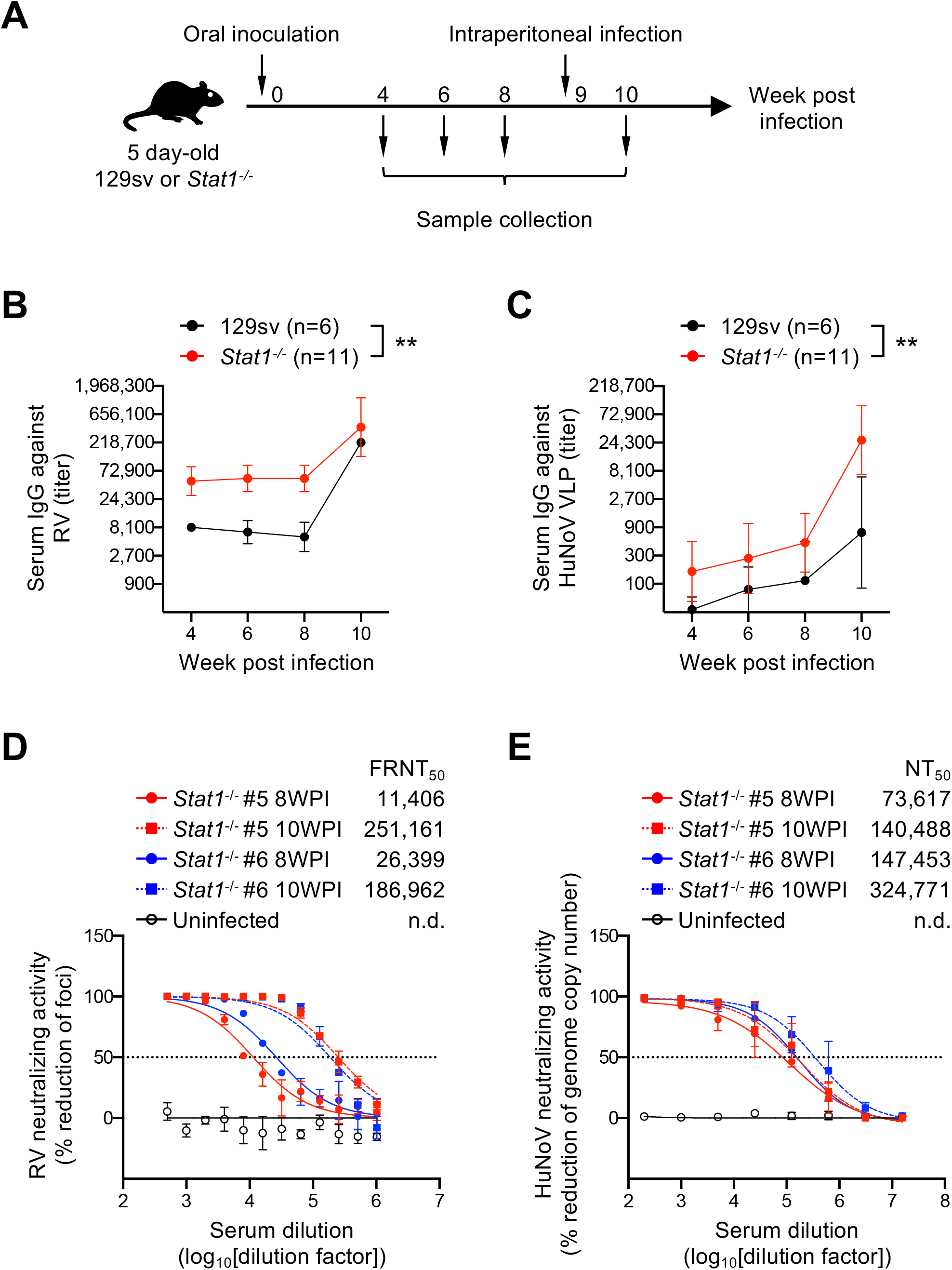
Systemic antibody responses against RV and HuNoV VLPs in 129sv and *Stat1*^*-/-*^ mouse following rRRV-HuNoV-VP1 infection. (**A**) Schematic presentation of the infection and immunization experiments. Five-day-old 129sv (n=6) or *Stat1*^*-/-*^ (n=11) pups were orally inoculated with rRRV-HuNoV-VP1 (3.9×10^5^ FFU/pup) at 0 WPI and intraperitoneally infected with the virus (3.9×10^5^ FFU/pup) at 9 WPI. Blood and stool samples were collected at 4, 6, 8, and 10 WPI. (**B and C**) Temporal dynamics of serum IgG titers against RV and HuNoV VLP. Mouse sera were serially diluted and tested in ELISA for (**B**) anti-RV IgG or (**C**) anti-HuNoV VLP. The IgG titer was defined as the highest serum dilution at which the OD score is higher than an uninfected mouse serum. Data are shown as mean with standard deviation. Statistical analysis was performed by two-way ANOVA. Statistical significance is indicated as **: P < 0.01. (**D**) Serum neutralization activities against RV. Mouse sera were serially diluted, mixed with RRV, and incubated at 37°C for 1 hour. The virus titer in the mixture of the serum and RRV was determined using MA104 cells in an FFU assay. Data are shown as mean scores of the percentage reduction of the focus number with standard deviation. (**E**) Serum neutralization activities against HuNoV. Mouse sera were serially diluted, mixed with human HuNoV GII.4 Sydney [P16] strain (GenBank # OL898515), and incubated at 37°C for 1 hour. The virus titer was determined using HIO monolayers and quantitative real-time RT-PCR 24 hours after inoculation. Data are shown as mean scores of the percentage reduction of the number of genomic copies.

To further characterize the serum immunoglobulin G (IgG) antibody responses induced in the 129sv and *Stat1*^*-/-*^ mice following rRRV-HuNoV-VP1 infection, we determined the ELISA binding titers of anti-RV and anti-HuNoV VLP IgG at different time points post inoculation. Consistent with the immunostaining data, all sera from both 129sv and *Stat1*^*-/-*^ mice infected with rRRV-HuNoV-VP1 reacted to RV as early as 4 WPI, and the serum anti-RV IgG titers increased further by 10 WPI following the IP boost (**Figure 3B and Table S1**). Remarkably, three of six 129sv mice also demonstrated seroconversion to HuNoV VLP at 6 WPI, showing a titer of 1:114 by 8 WPI (**Figure 3C**). The IP booster seroconverted two more mice that were negative at 8 WPI and enhanced the titer to 1:736 at 10 WPI (**Figure 3C and Table S1**). Sera from all *Stat1*^*-/-*^ mice showed higher serum anti-HuNoV VLP IgG titers than seen in the 129sv mice at all time points (1:494 at 8 WPI and 1:26,852 at 10 WPI) (**Figure 3C and Table S1**). Based on these findings, we concluded that sera from 129sv and *Stat1*^*-/-*^ infected with rRRV-HuNoV-VP1 contain antibodies against both RV and HuNoV VLP and that a parenteral IP boost enhanced the immune response to RV and HuNoV VP1.

### Neutralization of RV and HuNoV by sera from rRRV-HuNoV-VP1 infected *Stat1*^*-/-*^ mice

We further evaluated the neutralizing activity of serum antibodies against RV and HuNoV Because *Stat1*^*-/-*^ mice #5 and #6 showed the highest anti-HuNoV VLP binding titers among the eleven mice tested (**Table S1**), and because of limitations on our ability to carry out multiple HuNoV neutralization assays, we focused neutralization testing on specimens from these 2 mice (hereafter called sera #5 and #6) from 8 and 10 WPI. Not surprisingly, both sera #5 and #6 neutralized RV. Serum #5 had a 50% focus reduction neutralization titer (FRNT_50_) of 1:11,406 on 8 WPI and 1:251,161 on 10 WPI, while serum #6 showed a FRNT_50_ of 1:26,399 on 8 WPI and 1:186,962 on 10 WPI (**Figure 3D**). Human HuNoV recognizes the host HBGA molecules through the P2 subdomain in the P domain of the HuNoV VP1 protein (46-48). Expression of the P domain can form a self-assembling subunit P particle that exposes the P2 subdomain on its surface and binds to HBGA in the same manner as VLPs (42, 43, 49). As an alternative to a traditional restriction of replication-based neutralization assay, we first assessed whether sera #5 and #6 were capable of blocking P particle binding to HBGA, the cell surface receptor for HuNoV (43). Serum #5 inhibited P particle binding to HBGA at a 50% blocking titer (BT_50_) of 1:12.5 on 8 WPI and 1:50 on 10 WPI, while serum #6 inhibited binding at a BT_50_ of 1:6.25 on 10 WPI (**Table 1**). Furthermore, treatment of HuNoV with sera #5 or #6 prior to infection reduced the HuNoV genome copy number in a human intestinal organoid culture based neutralization assay (50). Serum #5 had a 50% neutralization titer (NT_50_) of 1:73,617 on 8 WPI and 1:140,488 on 10 WPI, while serum #6 had a NT_50_ of 1:147,453 on 8 WPI and 1:324,771 on 10 WPI (**Figure 3E**). Collectively, these data support the conclusions that rRRV-HuNoV-VP1 induced robust antibody responses against both RV and HuNoV in *Stat1*^*-/-*^ suckling mice and both post immunization sera developed the capacity to effectively neutralize HuNoV in two different neutralization assays.

**Table 1.** Blocking of HuNoV P particle binding to HBGA by sera from *Stat1*^*-/-*^ mice infected with rRRV-HuNoV-VP1.

| Sample | 50% blocking titer (BT<sub>50</sub>) of<br>HuNoV P particle binding to<br>HBGA |
| --- | --- |
| *Stat1*<sup>-/-</sup> #5 8WPI | 12.5 |
| *Stat1*<sup>-/-</sup> #5 10WPI | 50 |
| *Stat1*<sup>-/-</sup> #6 8WPI | <12.5 |
| *Stat1*<sup>-/-</sup> #6 10WPI | 6.25 |
| Uninfected | <3.125 |

### Induction of fecal IgA responses in 129sv and *Stat1*^*-/-*^ mice following rRRV-HuNoV-VP1 infection

To investigate whether rRRV-HuNoV-VP1 is capable of inducing a mucosal immune response in orally infected mice, we quantified the local enteric immunoglobulin A (IgA) responses to RV and HuNoV in stool specimens collected from the 129sv and *Stat1*^*-/-*^ immunized mice. Consistent with previous studies (28, 44, 51, 52), we detected RV-specific IgA in the fecal specimens from both the 129sv and *Stat1*^*-/-*^ immunized mice (**Figures 4A and 4B**). Of note, we found that the post infection fecal samples also contained HuNoV VLP-specific fecal IgA in the 129sv and *Stat1*^*-/-*^ mice (**Figures 4C and 4D**). Immunostaining with a secondary antibody specific to the α chain of murine IgA in *Stat1*^*-/-*^ mice also clearly demonstrated that the fecal samples contained HuNoV VP1 specific IgA (**Figure S5A**). Secretory IgA in the intestine has been frequently proposed to be a key component for the protection against enteric pathogens (53). Thus, we tested the ability of the fecal samples to neutralize both RV and HuNoV. We prepared fecal supernatants from *Stat1*^*-/-*^ mouse #2 and #4 (hereafter called fecal samples #2 and #4). These mice showed the highest ELISA values against HuNoV VLP at 10 WPI (**Figure 4D**). Fecal samples #2 and #4 demonstrated FRNT_50_ against RV at a dilution of 1/64% (w/v). A control stool suspension from uninfected mice showed a very low background reactivity at lower dilutions of 1/2 and 1/4% (w/v) (**Figure 4E**). Of note, fecal samples #2 and #4 also decreased the titer of HuNoV to the neutralization threshold in our human intestinal organoid assay using 1% (w/v) fecal suspensions (**Figure 4F**). These data suggest that rRRV-HuNoV-VP1 induced a functional enteric immune response against both RV and HuNoV in suckling mice.

**Figure 4.**
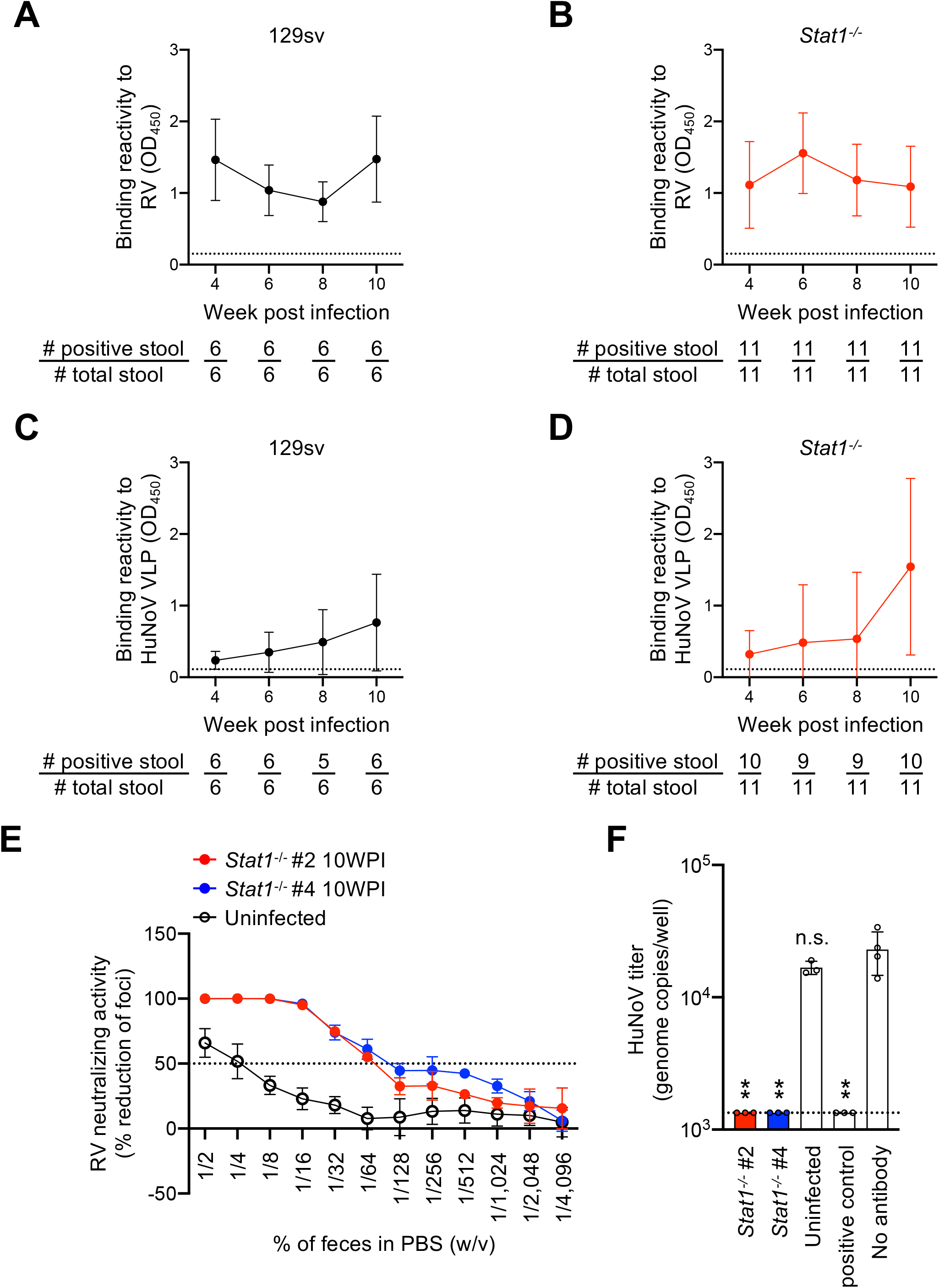
Mucosal antibody responses against RV and HuNoV VLPs in 129sv and *Stat1*^*-/-*^ mouse following rRRV-HuNoV-VP1 infection. (**A and B**) Temporal dynamics of fecal IgA responses to RV. The amount of fecal IgA against RV in (**A**) 1% (w/v) of fecal suspension from 129sv or (**B**) 0.1% (w/v) of fecal suspension from *Stat1*^*-/-*^ was determined by ELISA. Data are shown as the mean score of the net OD_450_ with standard deviation. The dotted lines show the limit of detection by stools from uninfected pups. A stool with a higher OD score than the dotted line was considered positive. The number of positive stools and total stools are shown below the graphs. (C and D) Temporal dynamics of fecal IgA responses to HuNoV VLP. The amount of fecal IgA against HuNoV VLP in 10% (w/v) of fecal suspension from (**C**) 129sv or (**D**) *Stat1*^*-/-*^ pups were determined by ELISA. Data are shown as the mean score of net OD_450_ with standard deviation. The dotted lines show background signals by stools from uninfected pups. A stool with a higher OD score than the dotted line was considered positive. The number of positive stools and total stools are shown below the graphs. (**E**) Fecal neutralization activities against RV. Mouse stool suspensions were serially diluted, mixed with RRV, and incubated at 37°C for 1 hour. The virus titer in the mixture of stool and RRV was determined using MA104 cells in an FFU assay. Data are shown as mean scores of the percentage focus reduction of the focus number with standard deviation. (**F**) Fecal neutralization activities against HuNoV. Mouse stool suspensions (1% [w/v]) were mixed with HuNoV GII.4 Sydney [P16] strain, incubated at 37°C for 1 hour, and inoculated to differentiated HIO monolayers. The virus titer in HIO was determined by quantitative real-time RT-PCR 24 hours after inoculation. Data are shown as mean scores of the HuNoV genome copy numbers. Statistical analysis was performed by one-way ANOVA test with Dunnett’s post-test. Statistical significance is indicated as ns: not significant, *: P < 0.5, **: P < 0.01.

## Discussion

In this study, we describe the generation and characterization of a recombinant RV expressing the HuNoV VP1 capsid protein that is capable of inducing both systemic and mucosal immune responses to both RV and HuNoV following oral administration of the recombinant RV to naïve suckling mice. Our data provide proof of principle that RV can be used as a “dual use” enteric vaccine vector (i.e., a possible live viral vaccine for combined RV and HuNoV immunization). We selected the RRV infection mouse model for these initial studies for several reasons. RRV has excellent *in vitro* replication characteristics as an expression vector. Insertion of exogenous genes into the RV genome often leads to a reduction in virus titer. Thus, it is preferable to use a RV that replicates to a high titer as a vaccine expression vector. In addition, inoculation of a high titer stock was likely more favorable for the enhanced expression of the exogenous protein target (in this case HuNoV VP1) *in vivo*. By virtue of the efficient *in vitro* replication capacity of RRV, rRRV-HuNoV-VP1 could be readily tested at a high dose in this initial proof of concept study. The majority of currently licensed RV vaccines are based either on human RVs or reassortants between between human and bovine RVs (11). It seems logical, if a vectored two-in-one RV-based enteric vaccine strategy is further pursued, to focus on modifying one of the already existing safe and effective licensed RV vaccines to express a second target antigen for use in humans as the most straightforward and practical path forward.

Of interest, RRV has already been extensively studied in both humans and animal models to better understand humoral and cellular immune responses to RV (26-28, 44, 45, 54-58). In fact, RRV represents the genetic backbone of a highly immunogenic human RV vaccine in multiple human clinical trials, but had issues related to safety in post-licensure surveillance which precluded its further use as a vaccine (29, 30). Prior studies of RRV infection in the suckling mouse model, including work by our group and others, documented enhanced RRV replication in mice deficient in components of innate immune signaling (26, 27, 45, 59-61). For the reasons above, we chose to study both immunocompetent 129sv and immunodeficient *Stat1*^*-/-*^ suckling mice. Of note, in a paper by Vancott (45), the antibody response against RV in *Stat1*^*-/-*^ mice was noted to be significantly higher than in 129sv mice following enteric RV infection. Given the reduced replication capacity of RRV in immunocompetent mice, we took advantage of the ability to enhance replication and immunogenicity in this only minimally permissive model system by using *Stat1*^*-/-*^ mice. As anticipated, rRRV-HuNoV-VP1 demonstrated enhanced replication in *Stat1*^*-/-*^ mice as assessed by the level of RV fecal shedding (**Figures 2B and 2D**) and by the induction of increased systemic and local intestinal origin antibody responses to RV and HuNoV VLP (**Figures 3C, 4C, and 4D**). These findings strongly suggest that there is a correlation between viral replication *in vivo* and the level of systemic and local antibodies in our study. Although *Stat1*^*-/-*^ mice are more permissive to RRV infection, the lack of STAT1 results in a defect in the interferon signaling pathway which could raise a concern that the immune response in *Stat1*^*-/-*^ mice may not reflect a natural immune response to rRRV-HuNoV-VP1 in an immunocompetent animal model. Fortuitously, we did not see a blunted humoral adaptive immune response to either RV or HuNoV in *Stat1*^*-/-*^ mice. Since young immunocompetent RV naïve rats have been shown to actually much more efficiently support RRV infection (58), we expect that follow-up immunization studies in the neonatal rat model will provide further information regarding the immune response to a vectored RV-based dual antigen vaccine constructs under more natural viral replication conditions.

To develop a HuNoV vaccine, several HuNoV VLPs have been assessed in clinical trials (39-41). In addition, some viral vectors expressing HuNoV VLPs have been used to induce a mucosal response against HuNoV VLPs in mice or humans (i.e., Venezuelan equine encephalitis virus replicon, vesicular stomatitis virus, and adenovirus) (5-8). Among these viruses, the adenovirus type 5-based HuNoV vaccine is the only one to have been studied in phase I human clinical trials (8, 40). In that model, development of immunity to the adenovirus type 5 expression vector itself would not add significant protection against a second important pediatric enteric infection and so RV immunity would need to be addressed by a separate vaccine.

By taking advantage of the natural intestinal tissue tropism of RV, we developed an RV-based viral vector against HuNoV that potentially could induce humoral immunity to two separate highly important pediatric enteric pathogens. rRRV-HuNoV-VP1 induced antibody responses not only in the systemic circulation but also in the GI tract where RVs and HuNoVs preferentially replicate (**Figures 3C, 4C, and 4D**). Importantly, we observed the induction of neutralizing antibodies in both sera and stool specimens in rRRV-HuNoV-VP1 inoculated *Stat1*^*-/-*^ mice despite the fact that the suckling mice are, at best, only semi-permissive for RRV replication (**Figures 3E and 4F**). Furthermore, we confirmed that the *Stat1*^*-/-*^ mouse sera blocked HuNoV P particle binding to HBGA (**Table 1**). Follow-up studies can examine whether the neutralization response to the HuNoV VP1 protein in *Stat1*^*-/-*^ mice (or possibly immunocompetent suckling rats) is similar or identical to that generated in humans after natural HuNoV infection.

Although rRRV-HuNoV-VP1 induced immune responses against HuNoV VLP, the HuNoV VP1 sequence (1,623 bp) was not stably maintained in the recombinant RRV (**Figures S1A and 2A**). Previous studies clearly demonstrated an association between the size of an exogenous gene and its genomic stability during RV replication. Recombinant RVs stably retained foreign genes during multiple rounds of replication when the gene size was less than 1 kbp (e.g., eGFP, ZsGreen, AsRed2, and mCherry [693∼720 bp], UnaG [420 bp], and Nano-luciferase [516 bp]) (12, 14-17, 19). On the other hand, recombinant simian SA11 strain with the gene for SARS-CoV-2 spike protein S1 fragment (2,055 bp) lost a part of the spike protein gene following 3 serial passages in cell culture (21). A potential solution to overcome the genetic stability of rRRV-HuNoV-VP1 is to replace the entire VP1 sequence with a smaller gene that expresses the HuNoV P domain (∼960 bp). The HuNoV P domain self-assembles into P particles when expressed with a tag sequence to assist P particle assembly (42). Administration of P particle has been shown to induce blocking antibody that inhibits HuNoV VLP binding to host HBGA receptors (43). Future studies to investigate the capacity of antibody induction by recombinant RV expressing HuNoV P particle are planned.

If these initial HuNoV immunization proof of concept studies prove successful, the RV-based dual vaccine vector system could be expanded to other small enteric immune target antigens. Potential candidates might include astrovirus capsid protein and *Escherichia coli* heat-labile enterotoxin produced by enterotoxigenic *Escherichia coli* (62, 63). The current findings provide an initial demonstration of the feasibility of developing a new approach to inducing mucosal immune responses to multiple enteric pathogens using a single enteric virus as simultaneously a live RV vaccine and a viral vaccine vector. The data presented here potentially have broad practical implications for the design of RV-based multivalent mucosal vaccine candidates against other common viral, bacterial, and parasitic enteric pathogens, ultimately reducing infectious diarrhea-associated morbidity and mortality worldwide.

## Materials and Methods

### Cells and Viruses

MA104 cells (ATCC CRL-2378.1) were cultured in Medium 199 (gibco, cat. #11150-059) supplemented with 10% Fetal Bovine Serum (FBS), 0.292 mg/ml of L-glutamine, 100 I.U./ml of penicillin, and 100 μg/ml of streptomycin (CORNING, cat. #30-009-CI). BHK-T7 cells were kindly gifted from Dr. Buchholz (NIH) (64), and cultured in DMEM (CORNING, cat. #10-013-CV) supplemented with 10% FBS, 0.292 mg/ml of L-glutamine, 100 I.U./ml of penicillin, and 100 μg/ml of streptomycin. Recombinant wild-type RRV was rescued as previously described and propagated in MA104 cells cultured in serum-free Medium 199 (SFM) in the presence of 0.5 μg/ml trypsin (Sigma-Aldrich, cat. #T0303) (12).

### Recombinant proteins and antibodies

Recombinant HuNoV VLP (MD145 strain) was generated as previously described (65). The following antibodies were used in this study: anti-RV double-layered particle rabbit antiserum generated in Greenberg’s lab, anti-RV triple-layered particle guinea pig antiserum generated in Greenberg’s lab, anti-HuNoV VLP guinea pig antiserum (403 anti-MD2004 virus [GII.4]) (66), anti-RV capsid protein mouse monoclonal antibody (Santa Cruz Biotechnology, clone 2B4, cat. #sc-101363), anti-β-actin mouse monoclonal antibody (Sigma-Aldrich, clone AC-74, cat. #A2228), HRP conjugated anti-mouse IgG goat polyclonal antibody (Sigma Aldrich, cat. #A4416), HRP conjugated anti-rabbit IgG goat polyclonal antibody (Sigma-Aldrich, cat. #A0545), HRP conjugated anti-guinea pig IgG goat polyclonal antibody (Sigma-Aldrich, cat. #A5545), Alexa Fluor488 conjugated anti-mouse IgG donkey polyclonal antibody (Jackson ImmunoResearch, cat. #715-545-150), Alexa Fluor488 conjugated anti-guinea pig IgG donkey polyclonal antibody (Jackson ImmunoResearch, cat. #706-545-148), Alexa Fluor594 conjugated anti-rabbit IgG donkey polyclonal antibody (Jackson ImmunoResearch, cat. #711-585-152). DyLight488 conjugated anti-mouse IgA goat polyclonal antibody (Abcam, cat. # ab97011). Rabbit antiserum to RV NSP3 was kindly gifted by Dr. Didier Poncet (67).

### Plasmid construction

The plasmid encoding African swine fever virus (ASFV) capping enzyme and T7 RNA polymerase (C3P3-G1) was kindly gifted by Dr. Philippe H. Jaïs (68). To construct the plasmid harboring HuNoV VP1 in RV gene 7 (pT7-RRV-NSP3-HuNoV-VP1), we amplified the gene (ORF2) encoding the VP1 sequence of the HuNoV Sydney strain by PrimeStar HS DNA polymerase (Takara Bio, cat. #r010a) and replaced it with the sequence encoding GFP in the pT7-RRV-NSP3-GFP constructed previously (12) by NEBuilder HiFi DNA Assembly Master Mix (New England Biolabs, cat. #E2621S). To construct expression vectors encoding HuNoV-VP1 (pCAG-HuNoV-VP1) and RV-VP6 (pCAG-RV-VP6), we cloned ORF2 of HuNoV Sydney strain VP1 or RV-VP6 in the pCAG vector by NEBuilder HiFi DNA Assembly Master Mix. The plasmid sequence was confirmed by DNA sequencing.

### Reverse genetics

Recombinant viruses were rescued by previously describe reverse genetics protocol with minor modification (12). Briefly, eight rescue plasmids (pT7-RRV-VP1, VP2, VP3, VP4, VP6, NSP1, NSP3, and NSP4) (0.4 μg of each), two rescue plasmids (pT7-RRV-NSP2 and NSP5) (1.2 μg of each) and C3P3-G1 (0.8 μg) were mixed with Trans-IT LT1 transfection Reagent (12.8 μl) (Mirus Bio, cat. #MIR2305) in OPTI-MEM I Reduced Serum Medium (150 μl) (Thermo Fisher Scientific, cat. #31985062). The mixture was transfected into BHK-T7 cells seeded in 12 well plates. The medium was replaced with serum-free DMEM 24 hours post-transfection. MA104 cells (1×10^5^ cells) were overlayed on the BHK-T7 cells 48 hours post-transfection and cultured for three days in the presence of 0.5 μg/ml of trypsin. The cells were frozen/thawed and passaged in MA104 cells. To rescue rRRV-HuNoV-VP1, we used pT7-RRV-NSP3-HuNoV-VP1 in place of pT7-RRV-NSP3. The rescued virus was propagated with MA104 cells in SFM in the presence of 0.5 μg/ml trypsin.

### RNA-PAGE analysis of viral dsRNA and genetic stability

rRRV-HuNoV-VP1 was passaged in MA104 cells by infection at an MOI of ∼1 FFU/cell. Viral dsRNAs were extracted from passages 3, 4, and 5 stocks of rRRV-HuNoV-VP1 with Trizol Reagent (Thermo Fisher Scientific, cat. #15596018) according to the manufacturer’s instruction. The extracted dsRNA was mixed with Gel Loading Dye Purple (6×) (New England Biolabs, cat. #B7024S) and separated by 10% polyacrylamide gel (25 mA at 4°C for 18 hours), then stained by silver staining.

### Growth kinetics

MA104 cells (seeded on 24 well plates) were infected with rRRV or rRRV-HuNoV-VP1 at MOI of 1 or 0.01 FFU/cell. One hour after inoculation, the inoculum was removed, and the cells were washed with the SFM twice and cultured with the SFM in the presence of 0.5 μg/ml of trypsin.. The cells were frozen at individual time points. Virus titer in the cells was determined by focus assay.

### Focus assay

Sample viruses were activated by 5 μg/ml of trypsin at 37°C for 15 minutes and serially diluted in SFM. MA104 cells (seeded on 96 well plates) were infected with the diluted samples. One hour after inoculation, the inoculum was removed and the cells were washed with SFM once, then cultured with SFM for 14 hours. The cells were fixed with 10% formalin (Fisherbrand, cat. #23-245685), permeabilized with PBS containing 0.05% Tween-20, and stained with rabbit anti-RV antiserum (dilution 1:1,000) at room temperature for 2 hours. The cells were stained with HRP conjugated anti-rabbit IgG at 4°C overnight. RV-positive cells were visualized by AEC Substrate Kit, Peroxidase (Vector Laboratories, cat. #SK-4200) and counted under a microscope. The virus titer was expressed as FFU/ml.

### Immunostaining

BHK-T7 cells (seeded on 96 well plates) were transfected with either pT7-RRV-NSP3, pCAG-HuNoV-VP1, or pT7-RRV-NSP3-HuNoV-VP1 (100 ng/well) by TransIT-LT1 and cultured for 36 hours. MA104 cells (seeded on 48 well plates) were infected with rRRV or rRRV-HuNoV-VP1 at an MOI of 1 FFU/cell and cultured for 24 hours or were transfected with pCAG-HuNoV-VP1 (250 ng/well) by TransIT-LT1 and culture for 3 days. The BHK-T7 cells and MA104 cells were fixed with 10% formaldehyde, permeabilized with PBS containing 0.1% triton-X, and blocked with PBS containing 2% FBS. The cells were stained with rabbit anti-RV NSP3 (dilution 1:1,000) and guinea pig 403 anti-HuNoV MD2004 virus (GII.4) VLP (dilution 1:1,000) at 37°C for 2 hours followed by Alexa488 conjugated anti-guinea pig IgG (dilution 1:1,000) and Alexa594 conjugated anti-rabbit IgG (dilution 1:1,000) at 37°C for 2 hours. The nuclei were stained with 4’,6-Diamidino-2-Phenylindole, Dilactate (Thermo Fisher Scientific, cat. #D3571). The images were acquired using an All-in-One Fluorescence Microscope BZ-X710 (Keyence).

### Western blotting

MA104 cells (seeded on 24 well plates) were infected with rRRV or rRRV-HuNoV-VP1 at an MOI of 4 FFU/cell and cultured for 12 hours. At 12 hours post-infection, the cells were lysed with Laemmli sample buffer (Bio-Rad, cat. #1610737) and boiled at 95°C for 5 minutes. The proteins were resolved with 4-15% Mini-PROTEAN TGX Gel (Bio-Rad, cat. #456-1086) and transferred to the PVDF membrane (Bio-Rad, cat. #162-0177) and blocked with PBS-T containing 3% Nonfat Dry Milk (Cell Signaling Technology, cat. #9999S) at 4°C overnight. The proteins were reacted with guinea pig 403 anti-HuNoV MD2004 virus (GII.4) VLP (dilution 1:1,000), rabbit anti-RV NSP3 (dilution 1:1,000), mouse anti-RV capsid protein (dilution 1:1,000), or anti-β-actin mouse monoclonal antibody (dilution 1:1,000) at room temperature for 2 hours, then reacted with appropriate HRP conjugated secondary antibodies. Protein expression was visualized with Supersignal West Femto Maximum Sensitivity Substrate (Thermo Scientific, cat. #34095) and acquired by ChemiDoc MP Imaging System (Bio-Rad).

### Mouse experiment

Wild type 129sv and *Stat1*^*-/-*^ mice were originally purchased from Taconic Biosciences Inc. and maintained at the Veterinary Medical Unit of the Palo Alto VA Health Care System. To assess diarrhea rate and fecal RV shedding, 5-day-old 129sv or *Stat1*^*-/-*^ pups were inoculated with rRRV or rRRV-HuNoV-VP1 (3.9×10^5^ FFU/pup) and monitored for diarrhea for 10 days. Stool samples were collected in 40 μl of PBS (+) (CORNING, Cat. #21-030-CV) and stored at −80°C until use. To evaluate serum and fecal antibody titers against RV and HuNoV VLPs, 5-day-old 129sv or *Stat1*^*-/-*^ pups were orally inoculated with rRRV-HuNoV-VP1 (3.9×10^5^ FFU/pup) and intraperitoneally injected with rRRV-HuNoV-VP1 (3.9×10^5^ FFU/pup) at 9 weeks post-inoculation (WPI). Blood and stool samples were collected at 4, 6, 8, and 10 WPI. Collected blood was centrifuged at 5,000 rpm for 5 minutes and the supernatant was used as serum samples. Stool samples were collected in pre-weighed 1.5 mL tubes and PBS (+) was added to make 10% (w/v) of fecal suspension and stored at −80°C until use. All animal experiment protocol is approved by Stanford Institutional Animal Care Committee.

### ELISA for detection of fecal RV antigens

RV fecal shedding was determined by sandwich ELISA. Briefly, ELISA plates (E&K Scientific Products, cat. #EK-25061) were coated with guinea pig anti-RV TLP antiserum (dilution 1:4,000), blocked with PBS containing 2% BSA (Sigma Aldrich, cat. #A7030) and incubated with RRV-infected MA104 cell lysate. After washing, 70 μl of PBS and 2 μl of fecal suspension were added to the plate and incubated at 4°C overnight. The amount of RV antigen was detected by rabbit anti-RV DLP antiserum (dilution 1:5,000) and peroxidase substrate (SeraCare, cat. #5120-0047). The OD450nm was detected by a microplate reader ELx800 (BIO-TEK).

### Immunostaining for detection of serum IgG or fecal IgA against RV and HuNoV VP1

BHK-T7 cells (seeded on 96 well plates) were transfected with pCAG-HuNoV-VP1 or pCAG-RV-VP6 (100 ng/well). Two days after transfection, the cells were fixed with 10% formaldehyde, permeabilized with PBS containing 0.1% Triton X-100, and blocked with PBS containing 2% FBS. The cells were stained with mouse sera (dilution 1:300) or fecal suspension (5% [w/v]) collected from virus-infected 129sv or *Stat1*^*-/-*^ mice and stained with Alexa488 anti-mouse IgG (dilution 1:1,000) or Alexa488 anti-mouse IgA (dilution 1:1,000). The images were acquired using an All-in-One Fluorescence Microscope BZ-X710 (Keyence).

### ELISA for detection of serum IgG and fecal IgA against RV and HuNoV VLP

To detect serum IgG and fecal IgA against RV, a sandwich ELISA was performed. ELISA plate was coated with rabbit anti-RV DLP antiserum (dilution 1:2,500), blocked with PBS containing 2% BSA, and incubated with RRV-infected MA104 cell lysate. After washing, the plates were incubated with serially diluted mouse sera or diluted fecal suspension (1% [w/v] for stools from 129sv, and 0.1% [w/v] for stools from *Stat1*^*-/-*^) at 4°C overnight. To detect serum IgG and fecal IgA against HuNoV VLP, direct ELISA was performed. ELISA plate was coated with HuNoV VLP (150 ng/well) and blocked with PBS containing 2% BSA. the plates were incubated with serially diluted mouse sera or 10% (w/v) of fecal suspension at 4°C overnight. The amount of serum IgG or fecal IgA was detected by HRP conjugated anti-mouse IgG (dilution 1:20,000) or HRP conjugated anti-mouse IgA (dilution 1:2,000) and visualized with peroxidase substrate. The OD450nm was detected by a microplate reader ELx800. For serum IgG time course, the titer was determined as the highest dilution at which OD score is higher than that from uninfected mouse serum.

### Blocking activities of mouse sera in the binding of HuNoV P particles to HBGA

Blocking activities of serum samples in the binding of HuNoV P particles to HBGA were determined by a saliva-based assay as previously described (43). Briefly, 96 well plates were coated with boiled saliva samples containing type B antigen and blocked with nonfat milk. Serially diluted mouse sera were mixed with P particle (GII.4 VA387 strain) (0.5 ng/μl) and added to the 96 well plates. The amount of bound P particle was detected by anti-HuNoV VLP rabbit antiserum (dilution 1: 3,300) followed by HRP conjugated anti-rabbit IgG. Fifty % blocking titer was determined as the highest dilution at which OD score is less than 50% of the OD score by P particle alone (no serum).

### RV neutralizing assay

Serum and fecal samples were serially diluted in SFM and mixed with 400 FFU/100 μl of RRV and incubated at 37°C for 1 hour. MA104 cells (seeded on 96 well plates) were infected with 100 μl of the mixture of RRV with serum or fecal suspension and incubated at 37°C for 1 hour. After one hour of incubation, the cells were washed with SFM and incubated with SFM for 14 hours. The cells were fixed with 10% formalin, permeabilized with PBS containing 0.05% Tween-20, and stained with rabbit anti-RV antiserum (dilution 1:1,000) at room temperature for 2 hours. The cells were stained with HRP conjugated anti-rabbit IgG (dilution 1:1,000) at 4°C overnight. RV-positive cells were visualized by AEC Substrate Kit, Peroxidase and counted under a microscope. RV neutralizing activities were determined as the percentage reduction in the number of foci compared to RRV alone (no serum).

### Human intestinal organoids (HIOs) cultures

Adult, secretor positive, HIO cultures (J2 cell line) derived from jejunal biopsies were grown and maintained as described previously (50). Briefly, HIOs were suspended in 20 µl of Matrigel™ (Corning, cat# 356231) and grown as undifferentiated 3D cultures in 500 µL IntestiCult™ Organoid Growth Medium (OGM) Human (STEMCELL™ Technologies, cat# 06010) supplemented with 10 µM Y-27632 (Sigma-Aldrich, cat# Y0503). Undifferentiated monolayers were produced by plating single cell suspensions obtained from 7 days old highly dense 3D cultures. Cells were resuspended in 100 μl of IntestiCult™ OGM Human with 10 µM Y-27632 and seeded in collagen IV (Sigma-Aldrich, cat#. C5533) pre-coated 96 well plates. After 24 hours at 37°C and 5% CO_2_, culture medium was replaced with differentiation medium [50/50; Base media / IntestiCult™ OGM Human Basal Medium (STEMCELL™ Technologies, cat# 100-0190)]. to induce cell differentiation. Cells were differentiated for 4 days. The base media compromises Advanced DMEM/F-12 (Gibco, cat# 12634010) supplemented with 1% GlutaMAX (Gibco, cat# 35050079), 1% Penicillin/Streptomycin (Gibco, cat# 15140122), and 1% 1M HEPES (Gibco, cat# 15630080).

### Human norovirus neutralizing assay

Serum and fecal samples were serially diluted in infection media (base media supplemented with 500 µM sodium glycochenodeoxycholate (GCDCA; Sigma-Aldrich cat# G0759) plus 50 µM ceramide (C2; Santa Cruz Biotechnology, cat# sc-201375A) and mixed with 100 TCID_50_/100 μl of GII.4 Sydney [P16] (GenBank # OL898515) and incubated at 37°C, for 1 hour. Differentiated HIE monolayers (seeded on 96 well plates) were infected with 100 μl of the mixture of HuNoV with serum or fecal suspension and incubated at 37°C, 5% CO_2_ for 1 hour. After that, monolayers were washed with base media and incubated with differentiation media supplemented with 500 µM GCDCA plus 50 µM C2 for 24 hours. Plates were frozen are −70°C. Viral RNA was extracted from cultures (cells and media) using MagMAX™ - 96 Viral RNA Isolation Kit (Applied Biosystems, cat# AMB18365) according to the manufacturer’s instructions. HuNoV RNA was detected by GI/GII TaqMan real-time RT-PCR (69). HuNoV neutralizing activities were determined as the percentage reduction in the number of genomic copies compared to HuNoV alone (no serum).

### Statistical analysis

Statistical analysis was performed using GraphPad Prism 8 (GraphPad Software, Inc.). One-way ANOVA was used for Figure 4E. Two-way ANOVA was used for Figures 1B, 1C, 2B, 2C 3B, and 3C. Neutralization titers (NT_50_) were determined from log-transformed non-linear, dose-response sigmoidal curve fit data for Figures 3D and 3E.

## Acknowledgments and funding sources

We thank members of Greenberg lab and Ding lab for the helpful discussion and Linda Jacob for her secretarial work. We thank Dr. Philippe H. Jaïs (Eukarÿs SAS) for sharing the C3P3-G1 plasmid, Dr. Susana López (Universidad Nacional Autónoma de México) for sharing the RRV rescue plasmid, Dr. Sean Tucker (Vaxart Inc.) for providing the plasmid encoding HuNoV VP1, and Dr. Didier Poncet (Université Paris-Saclay) for sharing rabbit antiserum to RV NSP3. The findings and conclusions in this article are those of the authors and do not necessarily represent the official position of the Centers for Disease Control and Prevention. This work is supported by Stanford Maternal and Child Health Research Institute Postdoctoral Support FY2020 (TK), R01 AI150796 (SD), R01 AI125249, U19 AI116484 NIH grants, and a VA Merit Grant (GRH0022) (HBG).

## Figure legends

**Figure S1. Strategy and validation of HuNoV VP1 expression from RV gene segment 7.**

(**A**) Schematic presentation of the modified gene segment 7. T2A peptide and HuNoV capsid protein VP1 were inserted before the stop codon of RRV NSP3. The numbers indicate the nucleotide length of each gene cassette and untranslated regions (UTRs). (**B**) Immunostaining analysis of protein expression by pT7-RRV-NSP3-HuNoV-VP1 in BHK-T7 cells. BHK-T7 cells were transfected with pT7-RRV-NSP3, pCAG-HuNoV-VP1, or pT7-RRV-NSP3-HuNoV-VP1 and fixed at 36 hours post transfection. The cells were stained with antibodies specific to RV-NSP3 (red), HuNoV-VP1 (green), and DAPI (blue). Scale bar: 25 μm. (**C**) Western blotting analysis of protein expression by pT7-RRV-NSP3 or pT7-RRV-NSP3-HuNoV-VP1 in BHK-T7 cells. BHK-T7 cells were transfected with pT7-RRV-NSP3 or pT7-RRV-NSP3-HuNoV-VP1 and harvested at 2 days post transfection. The cells were lysed with Laemmli buffer and resolved by SDS-PAGE. Protein expression of HuNoV-VP1, RV-NSP3, and β-actin was detected by the specific antibodies. The numbers show the molecular weights determined by the protein ladder.

**Figure S2. Detection of serum IgG against RV VP6 and HuNoV VP1 in sera from 129sv pups.**

(**A and B**) Immunostaining analysis with sera from 129sv pups inoculated with (**A**) rRRV-HuNoV-VP1 or (**B**) rRRV. BHK-T7 cells were transfected with pCAG-RV-VP6 or pCAG-HuNoV-VP1 and fixed at 2 days post transfection. The RV-VP6 and HuNoV-VP1 expressed in the cells were stained with indicated mouse serum and Alexa488 anti-mouse IgG. Scale bar: 100 μm. (**C**) Same as (**A and B**) except that the RV-VP6 and HuNoV-VP1 expressed in the cells were stained with control antisera and appropriate secondary antibodies. Scale bar: 100 μm.

**Figure S3. Detection of serum IgG against RV VP6 and HuNoV VP1 in sera from *Stat1***^***-/-***^ **pups.**

(**A and B**) Immunostaining analysis with sera from *Stat1*^*-/-*^ pups inoculated with (**A**) rRRV-HuNoV-VP1 or (**B**) rRRV. BHK-T7 cells were transfected with pCAG-RV-VP6 or pCAG-HuNoV-VP1 and fixed 2 days post transfection. The RV-VP6 and HuNoV-VP1 expressed in the cells were stained with each mouse serum and Alexa488 anti-mouse IgG. Scale bar: 100 μm. (**C**) Same as (**A and B**) except that the RV-VP6 and HuNoV-VP1 expressed in the cells were stained with control antisera and appropriate secondary antibodies. Scale bar: 100 μm.

**Figure S4. Detection of fecal IgA against RV VP6 and HuNoV VP1 in fecal specimens from 129sv pups**.(**A**) Immunostaining analysis with fecal supernatant from 129sv pups inoculated with rRRV-HuNoV-VP1. BHK-T7 cells were transfected with pCAG-RV-VP6 or pCAG-HuNoV-VP1 and fixed at 2 days post transfection. The RV-VP6 and HuNoV-VP1 expressed in the cells were stained with each mouse fecal supernatant and Alexa488 anti-mouse IgA. Scale bar: 100 μm. (**B**) Same as (**A**) except that the RV-VP6 and HuNoV-VP1 expressed in the cells were stained with control antisera and appropriate secondary antibodies. Scale bar: 100 μm.

**Figure S5. Detection of fecal IgA against RV VP6 and HuNoV VP1 in fecal specimens from *Stat1***^***-/-***^ **pups.**

(**A**) Immunostaining analysis with fecal supernatant from *Stat1*^*-/-*^ pups inoculated with rRRV-HuNoV-VP1. BHK-T7 cells were transfected with pCAG-RV-VP6 or pCAG-HuNoV-VP1 and fixed at 2 days post transfection. The RV-VP6 and HuNoV-VP1 expressed in the cells were stained with each mouse fecal supernatant and Alexa488 anti-mouse IgA. Scale bar: 100 μm. (**B**) Same as (**A**) except that the RV-VP6 and HuNoV-VP1 expressed in the cells were stained with control antisera and appropriate secondary antibodies. Scale bar: 100 μm.

**Table S1.**

**Serum IgG responses against RV VP6 and HuNoV VLP in individual 129sv and *Stat1***^***-/-***^ **mice following rRRV-HuNoV-VP1 inoculation.**

## Notes

### Competing Interest Statement

Harry B. Greenberg currently consults for the following companies: Pfizer, Vaxart, FluGen, and Aridis. The remaining authors disclose no conflicts.

