## Supplemental figure for "Mucosal and systemic neutralizing antibodies to norovirus and rotavirus by oral immunization with recombinant rotavirus in infant mice"

**Figure S1.** Strategy to express HuNoV VP1 from RV gene

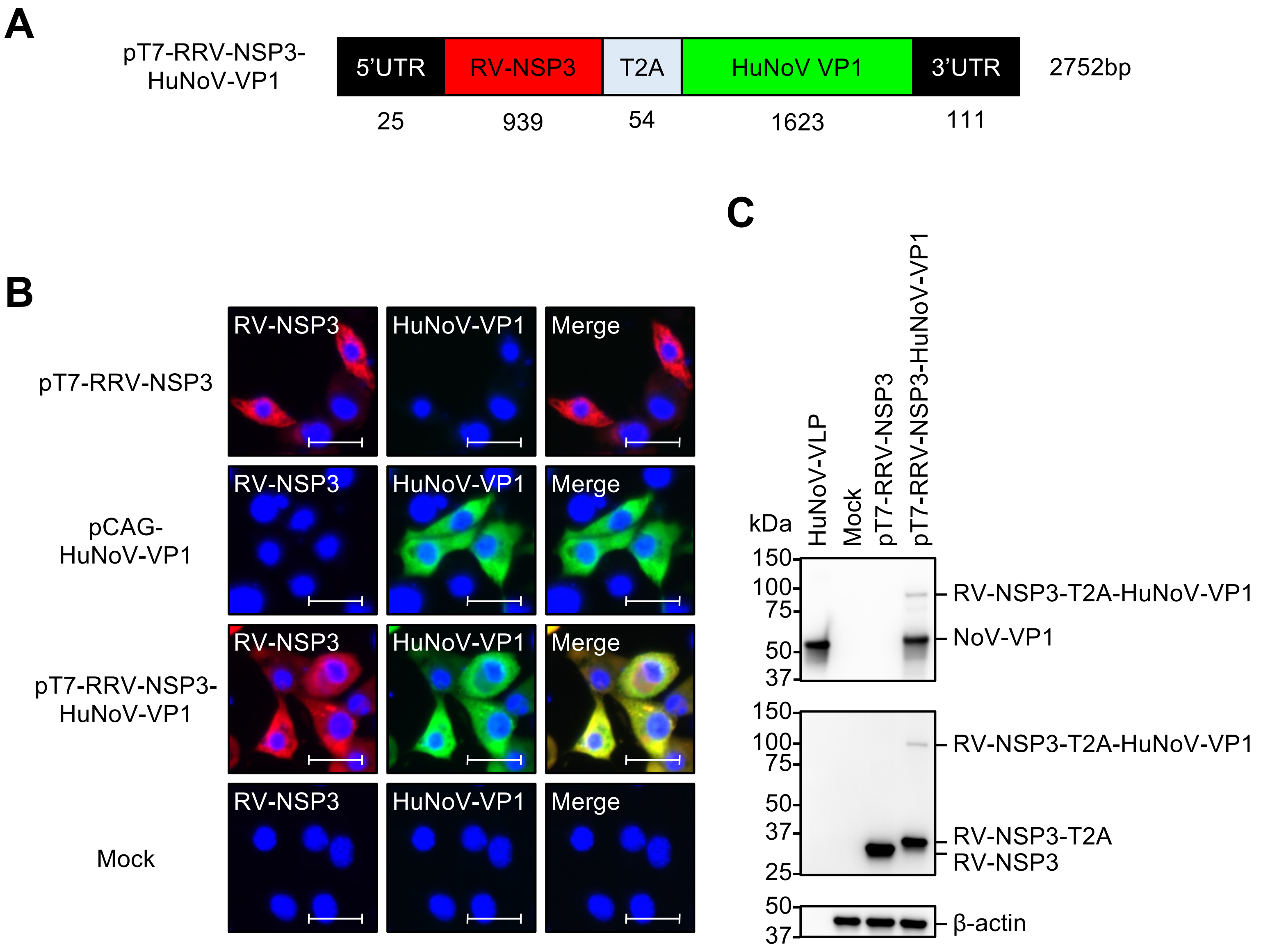

**Figure S2.** Detection of serum IgG against RV VP6 and HuNoV VP1 in sera from 129sv mice by immunostaining

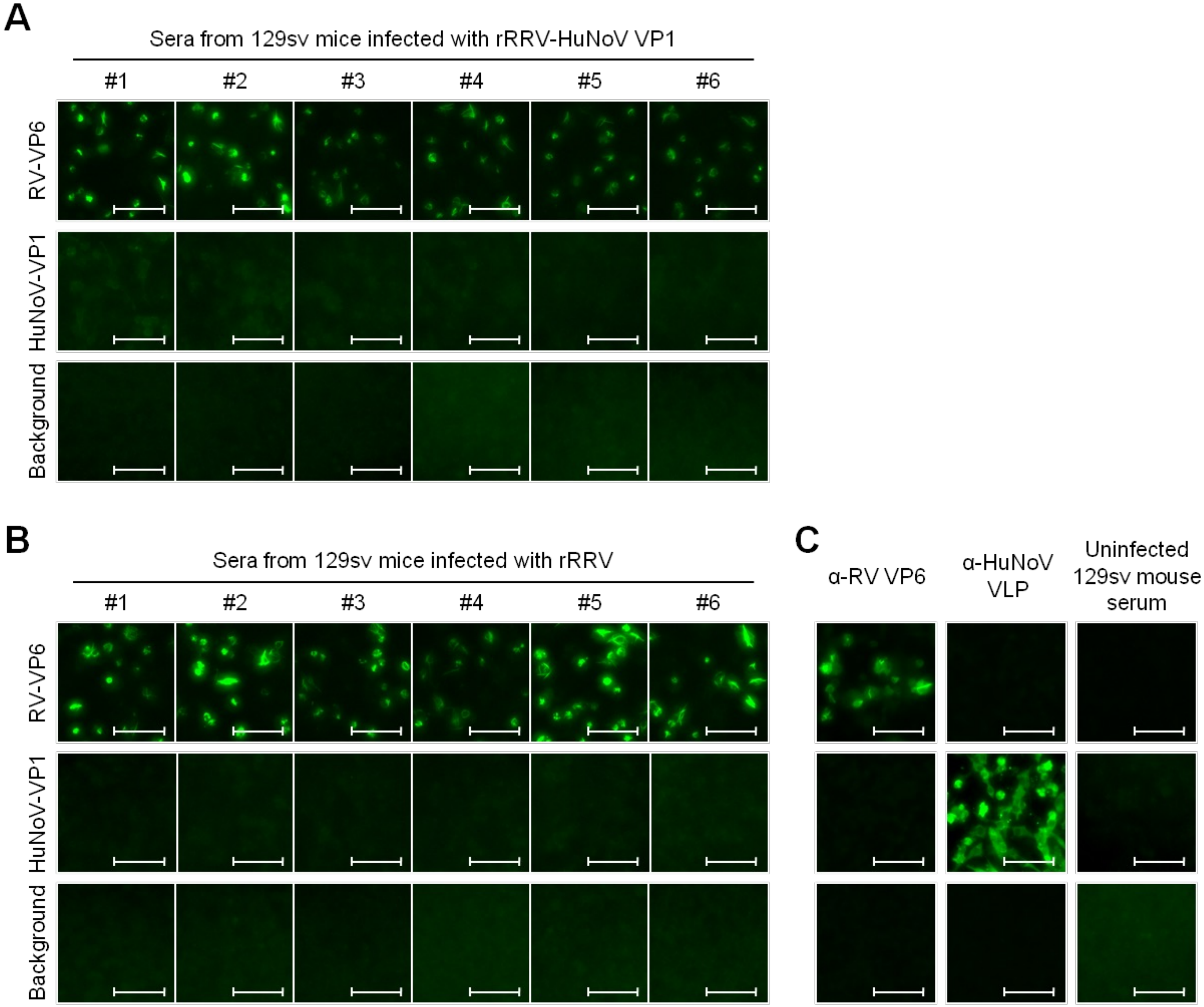

**Figure S3.** Detection of serum IgG against RV VP6 and HuNoV VP1 in sera from *Stat1*$^{-/-}$ mice by immunostaining

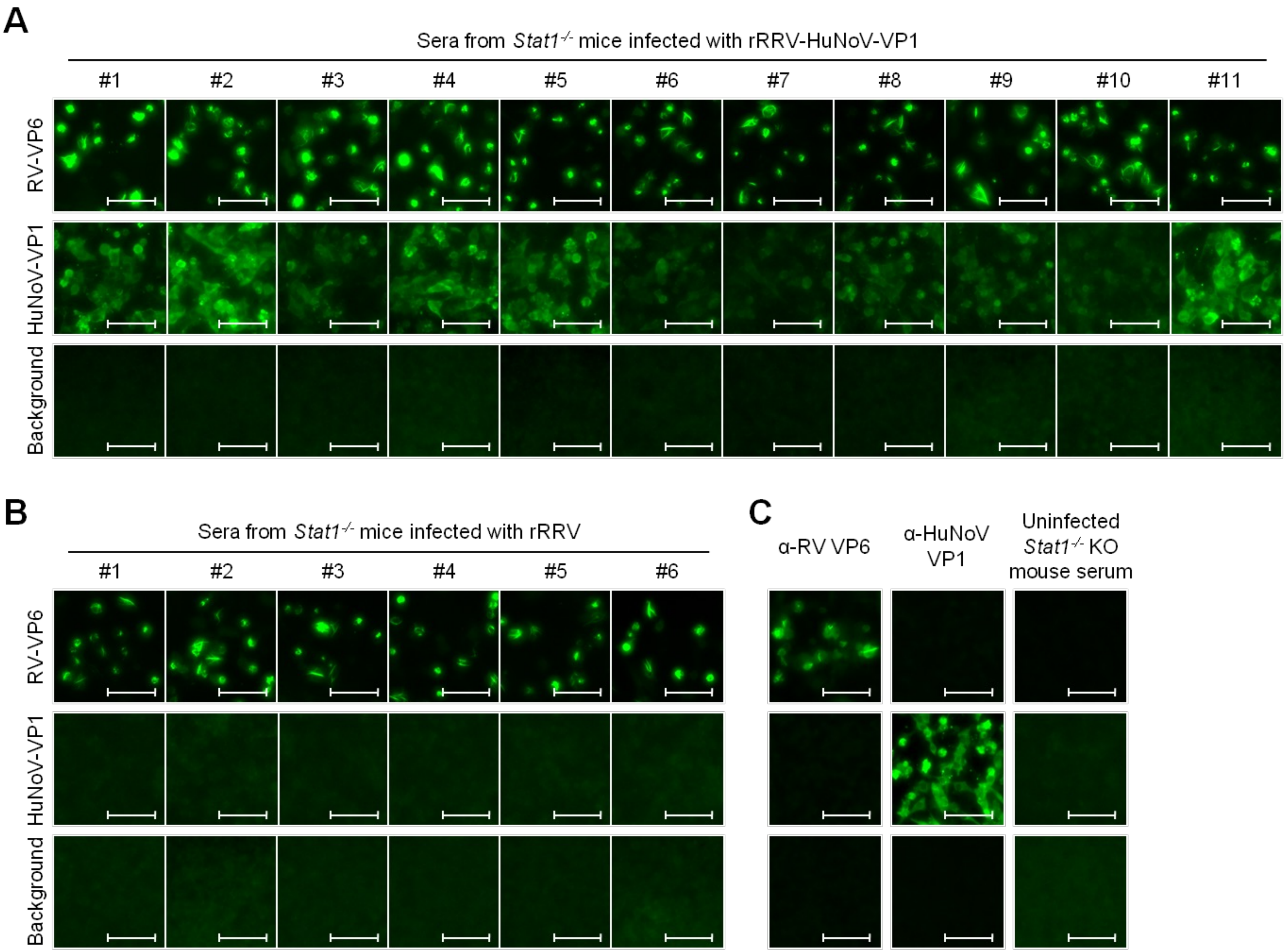

**Figure S4.** Detection of fecal IgA against RV VP6 and HuNoV VP1 in fecal supernatant from 129sv mice by immunostaining

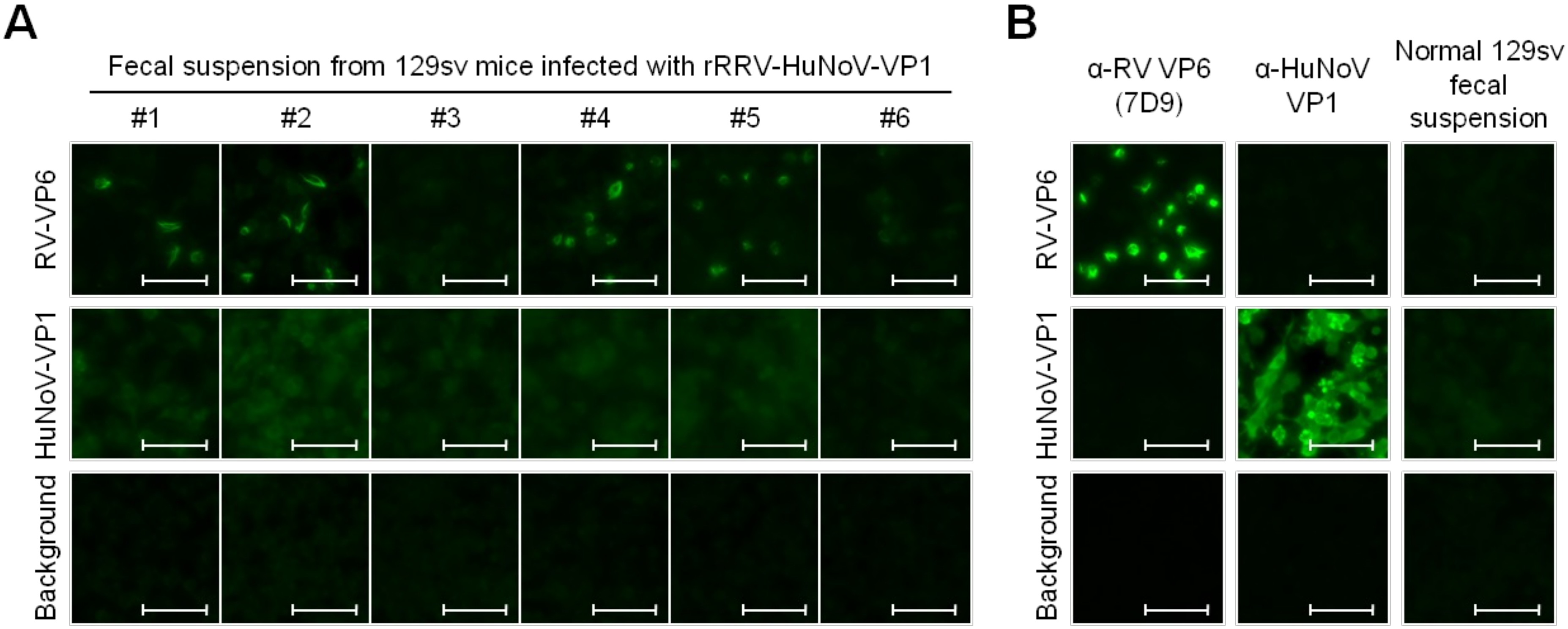

**Figure S5.** Detection of fecal IgA against RV VP6 and HuNoV VP1 in fecal supernatant from *Stat1*$^{-/-}$ mice by immunostaining

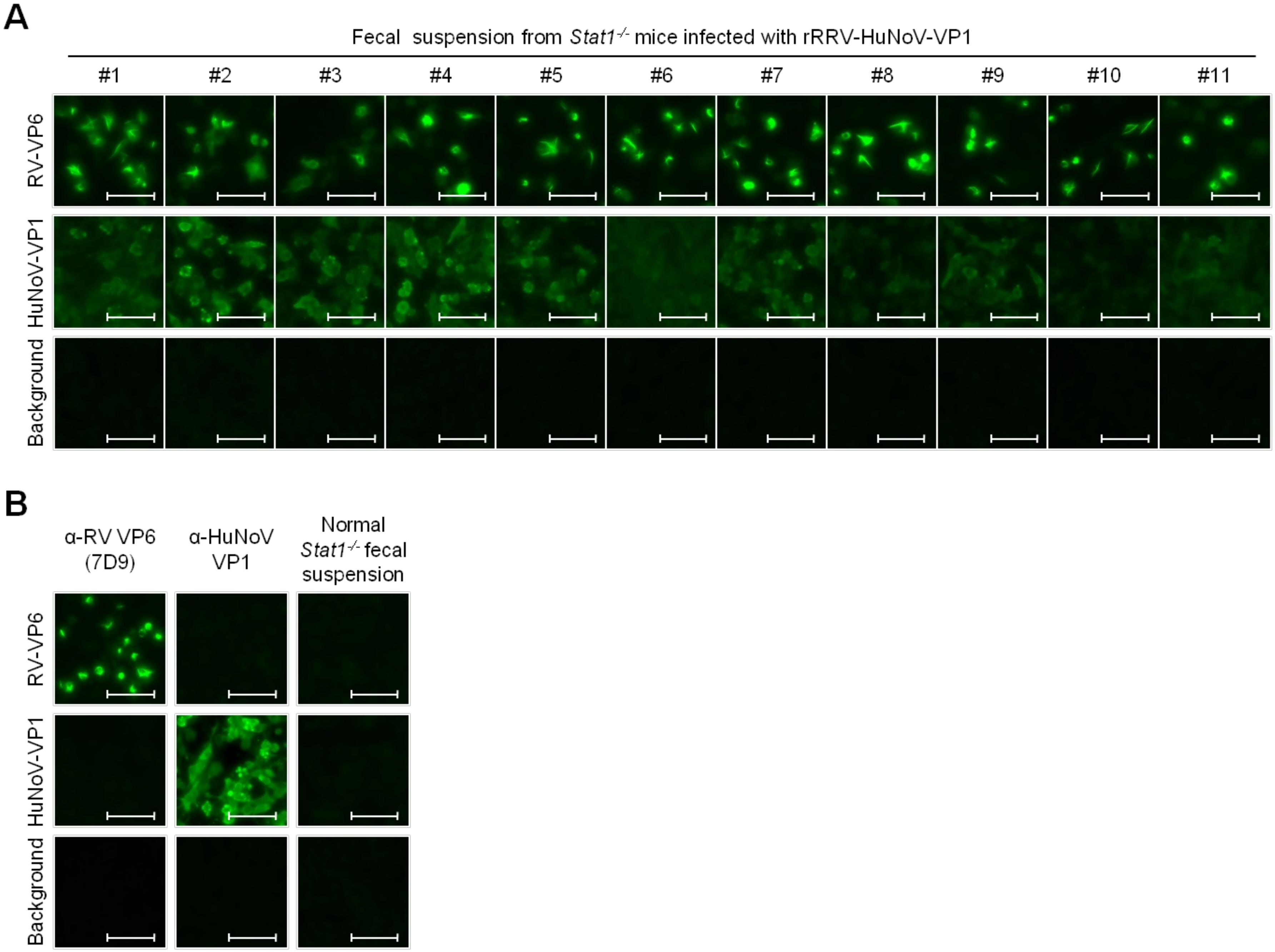

**Table S1.** Serum IgG responses against RV VP6 and HuNoV VLP in individual 129sv and *Stat1*$^{-/-}$ mice following rRRV-HuNoV-VP1 inoculation

| Animal ID | Serum IgG titer against RV | | Serum IgG titer against HuNoV | |
|---|---|---|---|---|
| | 8 WPI | 10 WPI (post booster shot) | 8 WPI | 10 WPI (post booster shot) |
| 129sv #1 | 8,100 | 218,700 | <100 | 300 |
| 129sv #2 | 2,700 | 218,700 | 300 | 900 |
| 129sv #3 | 8,100 | 218,700 | <100 | 900 |
| 129sv #4 | 8,100 | 218,700 | 300 | 24,300 |
| 129sv #5 | 8,100 | 218,700 | <100 | <100 |
| 129sv #6 | 2,700 | 218,700 | 900 | 900 |
| Average serum IgG titer (GMT) | 5,616 | 218,700 | 114 | 736 |
| % of serum IgG positive mice | 100 | 100 | 50 | 83 |
| *Stat1*$^{-/-}$ #1 | 72,900 | 656,100 | 300 | 24,300 |
| *Stat1*$^{-/-}$ #2 | 24,300 | 218,700 | 900 | 72,900 |
| *Stat1*$^{-/-}$ #3 | 24,300 | 72,900 | 100 | 2,700 |
| *Stat1*$^{-/-}$ #4 | 72,900 | 72,900 | 900 | 2,700 |
| *Stat1*$^{-/-}$ #5 | 72,900 | 656,100 | 2,700 | 72,900 |
| *Stat1*$^{-/-}$ #6 | 72,900 | 656,100 | 2,700 | 72,900 |
| *Stat1*$^{-/-}$ #7 | 72,900 | 1,968,300 | 300 | 72,900 |
| *Stat1*$^{-/-}$ #8 | 24,300 | 218,700 | 300 | 72,900 |
| *Stat1*$^{-/-}$ #9 | 72,900 | 656,100 | 300 | 8,100 |
| *Stat1*$^{-/-}$ #10 | 72,900 | 218,700 | 100 | 24,300 |
| *Stat1*$^{-/-}$ #11 | 72,900 | 1,968,300 | 900 | 72,900 |
| Average serum IgG titer (GMT) | 54,026 | 398,196 | 494 | 26,852 |
| % of serum IgG positive mice | 100 | 100 | 100 | 100 |
